# A comprehensive toolkit to enable MinION sequencing in any laboratory

**DOI:** 10.1101/289579

**Authors:** Miriam Schalamun, David Kainer, Eleanor Beavan, Ramawatar Nagar, David Eccles, John P. Rathjen, Robert Lanfear, Benjamin Schwessinger

**Author notes:** Corresponding authors: Robert Lanfear, Benjamin Schwessinger.

## Abstract

Long-read sequencing technologies are transforming our ability to assemble highly complex genomes. Realising their full potential relies crucially on extracting high quality, high molecular weight (HMW) DNA from the organisms of interest. This is especially the case for the portable MinION sequencer which potentiates all laboratories to undertake their own genome sequencing projects, due to its low entry cost and minimal spatial footprint. One challenge of the MinION is that each group has to independently establish effective protocols for using the instrument, which can be time consuming and costly. Here we present a workflow and protocols that enabled us to establish MinION sequencing in our own laboratories, based on optimising DNA extractions from a challenging plant tissue as a case study. Following the workflow illustrated we were able to reliably and repeatedly obtain > 8.5 Gb of long read sequencing data with a mean read length of 13 kb and an N50 of 26 kb. Our protocols are open-source and can be performed in any laboratory without special equipment. We also illustrate some more elaborate workflows which can increase mean and average read lengths if this is desired. We envision that our workflow for establishing MinION sequencing, including the illustration of potential pitfalls, will be useful to others who plan to establish long-read sequencing in their own laboratories.

## Introduction

Single-molecule nanopore sequencing records changes in electrical current as individual tagged DNA molecules pass through an engineered pore across a chemical gradient (Jain, Olsen, Paten, & Akeson, 2016). Groups of consecutive bases cause a characteristic shift in current, and this can be deconvoluted to infer the individual base sequence of the DNA molecule, a process referred to as basecalling. This technology can sequence DNA fragments of varied lengths, from a few hundred bases to over a megabase (Mb), which compares favorably to sequencing by synthesis (e.g. Illumina), which is limited to hundreds of bases (Leggett & Clark, 2017). Long reads have a number of important applications, including: improving the accuracy and efficiency of genome assembly, especially for genomes that contain long low-complexity regions; detailed investigation of segmental duplications and structural variation (Jain et al., 2018); major histocompatibility complex (MHC) typing (Liu et al., 2017); and detecting methylation patterns (Simpson et al., 2017). The number of genome assemblies using nanopore data either exclusively or in combination with other sequencing data is steadily increasing, for example the 3.5 gigabase (Gb) human genome, the 860 Mb European eel genome, the 1 Gb genome of the wild tomato species *Solanum pennellii*, and the 135 Mb genome of *Arabidopsis thaliana* (Jain et al., 2018; Jansen et al., 2017; Michael et al., 2018; Schmidt et al., 2017). In short, nanopore sequencing solves the technical challenges of reading long DNA fragments, while still having room for improvement in terms of accuracy. One of the primary remaining challenges is to extract and purify very long DNA fragments from the organisms or tissues of interest.

The Oxford Nanopore Technologies (ONT) MinION makes long-read sequencing accessible to most laboratories outside of a dedicated genome facility. It has very low capital cost; has the potential to generate more than 1 Gb of sequence data per 100 USD; has a footprint about the size of an office stapler; and runs on a standard desktop or laptop computer. The MinION uses small consumable flowcells for sequencing, which contain fluid channels that flow samples onto a sequencing matrix, and provide a small amount of fluid waste storage.

This democratization of sequencing brings the challenge that every laboratory has to establish the sequencing platform and concomitantly, new DNA extraction and library preparation protocols. This can be challenging and time consuming. Here we illustrate the workflow we applied to establish MinION sequencing in our laboratories using the tree species *Eucalyptus pauciflora* as a case study. It is challenging to extract high purity and high molecular weight DNA from *E. pauciflora* because the mature leaf tissue is physically tough, and because it contains very high levels of secondary metabolites which are known to reduce the efficacy of DNA extraction protocols (Healey, Furtado, Cooper, & Henry, 2014). We illustrate reliable and repeatable ways of measuring DNA purity to optimise output from the MinION sequencer. We discuss important considerations for DNA library preparation, and methods to control and optimise the final distribution of read lengths. We show that during DNA extraction, small alterations in sample homogenisation protocols can drastically alter DNA fragment lengths; introduce a novel low-tech size selection protocol based on Solid Phase Reversible Immobilization (SPRI) beads; and compare size selection methods using electrophoresis versus DNA shearing. Finally, we introduce an open-source MinION user group that shares DNA extraction, size-selection, and library preparation protocols for many additional organisms, making our workflow applicable well beyond the case study presented here.

## Results

### Optimizing sequencing output

#### DNA Sample Purity

The first goal of our project was to optimize extraction protocols to yield highly intact and high purity DNA suitable for long-read sequencing. High purity of DNA is defined by Nanodrop spectrophotometer (Thermo Fisher) absorbance of DNA with a 260/280 nm ratio between 1.8 to 2.0, and a 260/230 nm ratio between 2 and 2.2 (Desjardins & Conklin, 2010; Mackey & Chomczynski, 1997). In addition to this we found it critical that the ratio of DNA concentrations measured on the Qubit and Nanodrop instruments respectively should be 1:1.5. This ratio indicates that most DNA molecules are double-stranded and that no other molecules (e.g. RNA) are present that absorb at 260 nm (e.g. Qubit: 100 ng/µl; Nanodrop: 150 ng/µl gives an acceptable ratio of 1:1.5; (O’Neill, McPartlin, Arthure, Riedel, & McMillan, 2011).

In our workflow, we first aimed to recover high molecular weight DNA with a Nanodrop/Qubit concentration ratio that was close to one. We then optimized DNA purity based on 260/280 nm ratios, which are indicative of protein contamination, and 260/230 nm ratios, which are indicative of contamination by salts, phenol, and carbohydrates (O’Neill et al., 2011). To achieve this, we first tested a well-established hexadecyltrimethylammonium bromide (CTAB) extraction protocol to extract DNA from *E. pauciflora* leaves collected in June 2017 from adult trees in the Kosciuszko National park near Thredbo, New South Wales, Australia (Healey et al., 2014; Schwessinger & Rathjen, 2017). While the CTAB protocol returned good yields of double stranded DNA (∼5 µg DNA per g tissue), the Qubit/Nanodrop ratio of 0.05 indicated significant contamination with RNA or single-stranded DNA. Nanodrop absorption spectra from 220 to 350 nm (Figure 1A) revealed the presence of contaminants as the curve deviated drastically from pure DNA absorption curves (Figure 1D). In such cases it is often recommended to clean the DNA using SPRI paramagnetic beads in combination with a polyethylene glycol (PEG) and sodium chloride (NaCl) mixture, such as AMPure XP beads (Beckman Coulter). These beads bind to the DNA but most contaminants do not and can be washed away (Krinitsina, Sizova, Zaika, Speranskaya, & Sukhorukov, 2015; Mayjonade et al., 2016). We were able to improve sample quality slightly by adding the standard measure of 0.45 vol (V/V) AMPure XP beads (Figure 1B), but repeating this step did not increase the purity of the DNA further. Next we tested an extraction method employing the detergent sodium dodecyl sulfate (SDS) which contains a PEG-NaCl precipitation step to capture the DNA onto SPRI beads. This approach has been reported to work well with many species including sunflower, human, and *Escherichia coli* (Mayjonade et al., 2016). Using this approach we recovered high levels of double-stranded DNA (Qubit/Nanodrop = 1:1.5), but the Nanodrop absorption curves still indicated the presence of contaminants in the final DNA extract (Figure 1C). Again, we were unable to improve the DNA purity by repeated SPRI clean-up steps as the 260/280 nm ratios did not improve. As an alternative method we cleaned the crude DNA obtained from the SDS-based method using a chloroform:isoamylalcohol extraction followed by isopropanol precipitation of the DNA, as described for some fungal DNA samples (Dong, 2017). This consistently resulted in high purity DNA with Qubit/Nanodrop ratios of 1:1-1.5, 260/280 nm ratios of ∼1.8, 260/230 nm ratios of ∼2.0, and excellent Nanodrop absorbance curves (Figure 1D).

**Figure 1:**
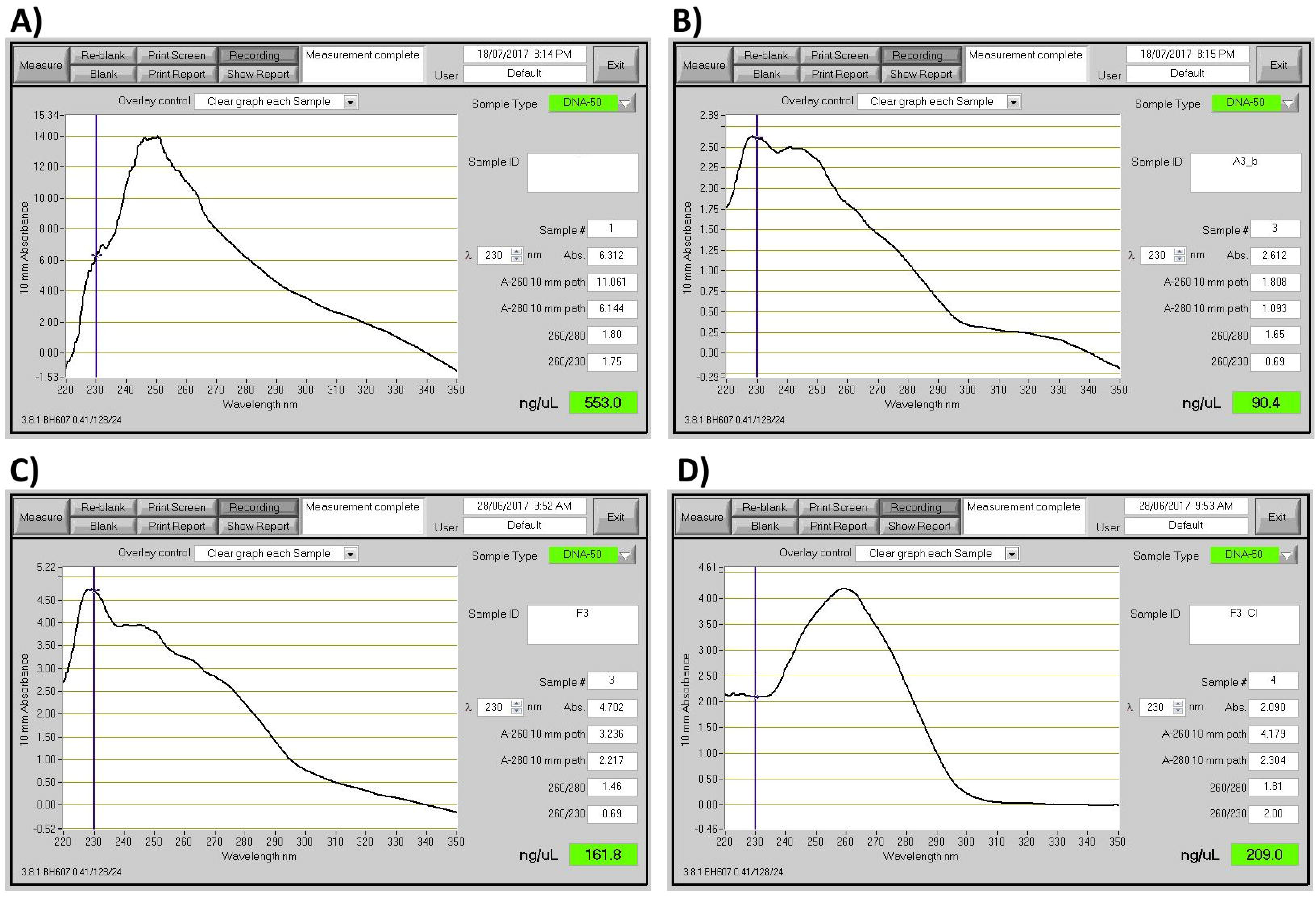
Illustration of different purity DNA preparations. Nanodrop readings of different DNA preparations. (A) DNA extraction with CTAB lysis buffer followed by phenol:chloroform:isoamylalcohol extraction (Schwessinger & Rathjen, 2017). (B) Sample A after SPRI beads clean-up. (C) DNA extraction using SDS lysis buffer and SPRI beads purification (Mayjonade et al., 2016). (D) Sample C followed by an additional chloroform:isoamylalcohol purification step.

ONT 1D library preparations involve the ligation of sequencing adapters at both 3’ ends of end-repaired double-stranded DNA. Sequencing adapters carry a motor protein that guides the DNA to the pore and regulates the translocation speed of the DNA across the pore. In addition, they carry a characteristic DNA sequence which is used by basecallers to recognize the translocation start of a new DNA molecule (Jain et al., 2016; Leggett & Clark, 2017). We tested the effect of sample impurities on MinION output using the 1D ligation protocol. Our three samples differed primarily in their 260/230 nm ratios. One sub-optimal sample (sample 5, Table 1) had a low ratio of 1.0, and the other two samples (samples 10 and 27, Table 1) had close-to-optimal ratios of 2.1 and 2.3 respectively. The sample with the low 260/230 nm ratio yielded an order of magnitude less sequence data from a single flowcell compared to the other two samples (0.7 Gb vs. ∼7 Gb respectively, Table 1, Supplemental Table 1). It seems likely that the contaminants causing the reduced 260/230 nm ratio inhibited the library preparation or the sequencing itself. Based on this observation we recommend adhering to the DNA quality measures nominated above whenever possible, or else to assume reduced sequencing outputs. We also advise establishing suitable DNA extraction methods well in advance of ordering sequencing materials; our experience suggests that optimizing DNA extraction protocols can take several months.

**Table 1.** DNA purity impacts sequencing yields. Comparison of yield per flowcell for different quality samples. Impact of sample quality measured by 260/280 and 260/230 nm ratios (Nanodrop data) on the final sequence output measured in Gb per flowcell (Figure 1). Sample #10 and #27 are two representative sequencing runs. #5 is a run with low input DNA purity.

| Sample | Qubit [ng/μl] | Nanodrop [ng/μl] | 260/280 | 260/230 | Yield [Gb] | Yield<sub>Q7</sub> [Gb] |
| --- | --- | --- | --- | --- | --- | --- |
| 10 | 178 | 203 | 1.8 | 2.1 | 6.0 | 5.9 |
| 27 | 142 | 188 | 1.8 | 2.3 | 7.8 | 7.4 |
| 5 | 57 | 80 | 1.7 | 1.0 | 0.7 | 0.7 |

#### Sequencing library preparation

The manufacturer-recommended library preparations involving DNA repair and end-prep are optimized for 0.2 pmol of input DNA with an average fragment size of 8 kb, which in turn requires 1 µg of double-stranded DNA. This implies that the DNA input as expressed in mass needs to be adapted according to the concentration of free DNA ends available for adapter ligation, which is a function of fragment length (Mayjonade, 2018; Schwessinger, 2018). The molarity of the DNA sample can be calculated using the Promega BioMath calculator (http://www.promega.com/a/apps/biomath/) which requires the average fragment length to calculate the respective DNA mass for 0.2 pmol. For example, 0.2 pmol of DNA of mean length 24 kb requires a DNA input of 3 µg. In our case we estimated a mean DNA fragment length of ∼30 kb based on 1% agarose gel electrophoresis (Figure 2) and pulsed field gel electrophoresis (PFGE) which provides higher resolution in the high molecular weight range (Figure 3). When estimating mean DNA fragment length based on fluorescent intensity (e.g. after staining with SYBR red or ethidium bromide) it is important to consider that smaller DNA molecules incorporate less dye so appear fainter during imaging. For example, even faint DNA smears below 10 kb can indicate the significant presence of short DNA fragments that are best avoided if long-read lengths are a primary goal of the sequencing effort (see below.) Failure to account for this can easily lead to overestimation of mean DNA fragment length, and miscalculation of the true concentration of DNA fragments.

**Figure 2.**
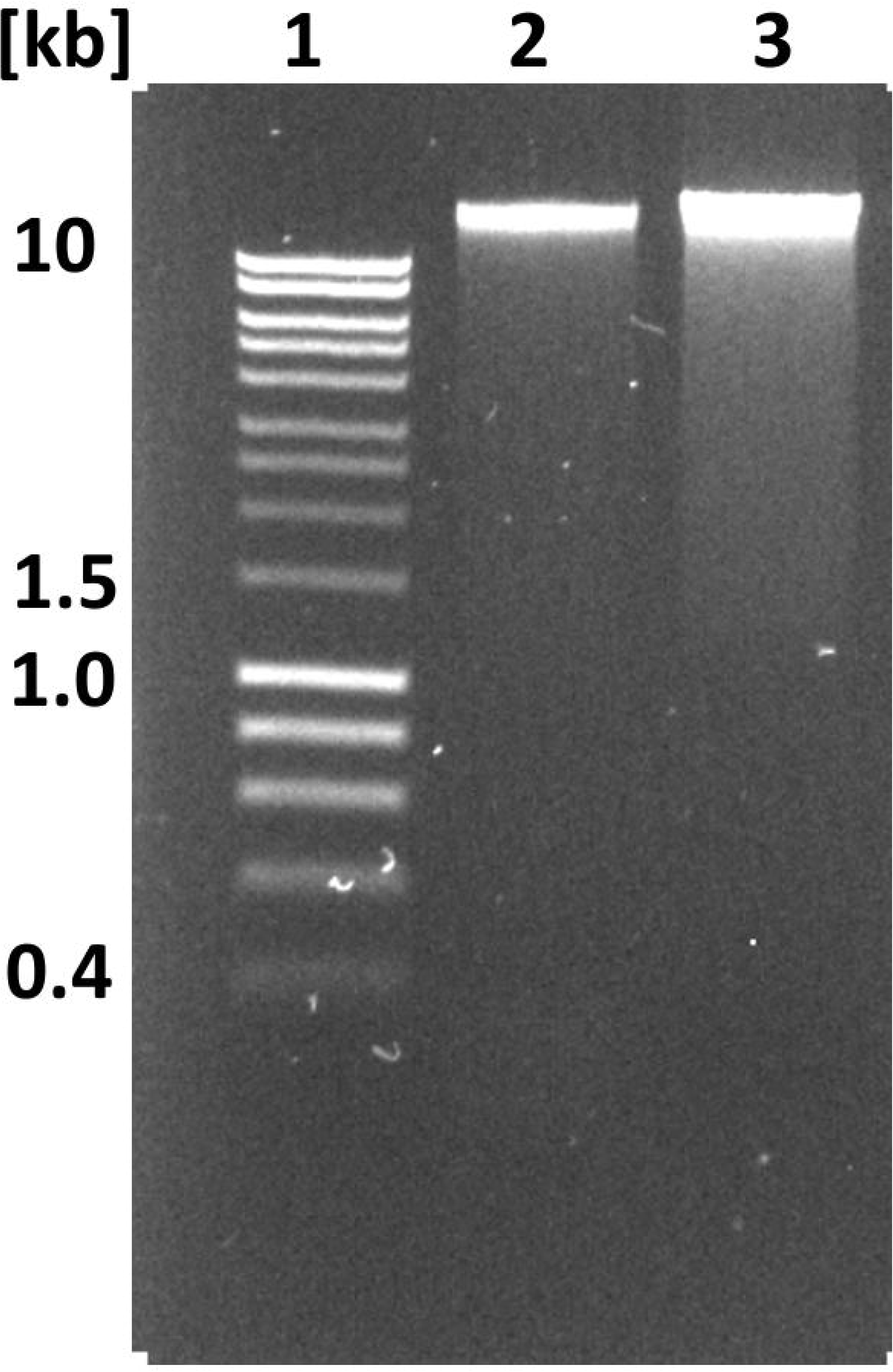
Illustration of the impact on DNA extraction procedures on DNA fragment length. 0.8 % agarose gel of 100 ng DNA prepared with two different DNA extraction procedures as explained in the main text. #1 HyperLadder 1 kb (Bioline). #2 DNA extracted following the default HMW DNA extraction protocol with mean read length of 13 kb as shown in Table 2. #3 DNA accidentally sheared during the extraction procedure with mean read length of 5 kb as shown in Table 2.

**Figure 3.**
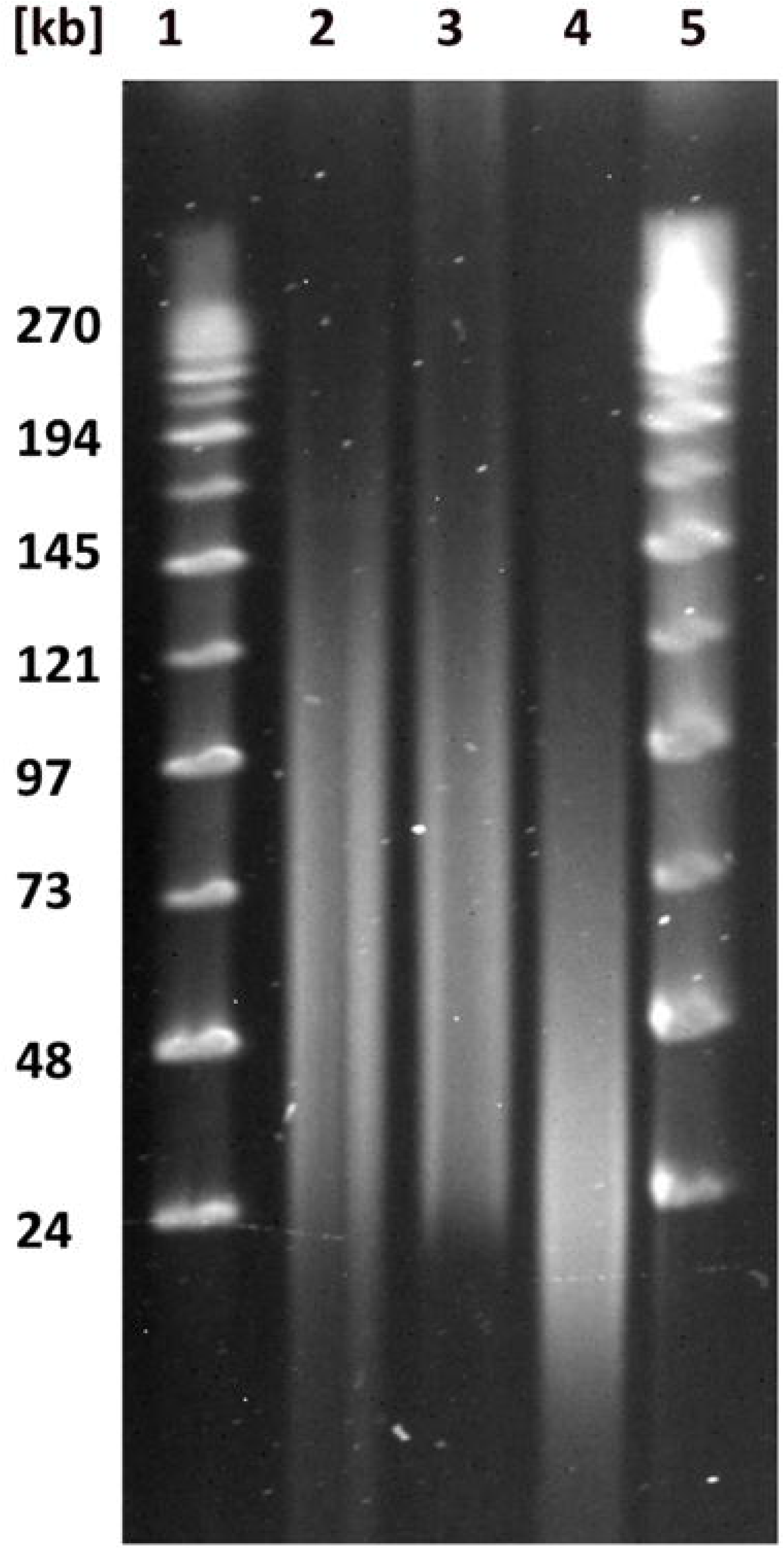
Purposeful mechanical shearing and high-pass filtering alters DNA fragment length distribution. Pulsed field gel electrophoresis of differently treated DNA samples. Lane #1 and #5 MidRange II PFG marker (BioLabs). Lane #2 DNA extracted following the default HMW DNA extraction protocol (mean read length of 13 kb as shown in Table 4). Lane #3 same DNA extraction as in #2 followed by size selection with the Blue Pippin using 20 kb high pass filtering (a mean read length of 26 kb as shown in Table 4). Lane #4 same same DNA extraction as in #2 followed by mechanical shearing with the a g-TUBE (a mean read length of 11.8 kb as shown in Table 3).

As a starting point we defined the optimal DNA input based on our initial mean fragment length estimate of 30 kb. This was followed by empirical adjustments from plotting sequencing outputs vs. the DNA input into adapter ligation (Figure 4). This approach revealed an optimum of ∼2 µg dsDNA (Figure 4), which required an input of 2.9 µg DNA for the DNA preparation stage considering typical losses of 30% after clean-up using in house SPRI beads (see below). Neither decreasing or increasing the DNA input improved the sequencing output, due to too few adapter-DNA molecules, or too many free DNA molecules damaging DNA pores. Assuming that 2.9 µg input DNA was the equivalent of 0.2 pmol, we estimate a mean DNA fragment length of 23 kb for our sample preparation. This suggests we initially overestimated the mean DNA fragment length, and highlights the difficulty of estimating these values based on gel imaging. At the same time, we stress that it is best to establish optimal DNA inputs empirically for each DNA extraction and/or shearing protocol. In addition, one can use the sequence read-length distribution from the initial flowcells to improve the estimate of the mean fragment length of the DNA extracted from the tissue. This approach can help to quickly optimise the amount of input DNA added to the ligation step.

**Table 2.** DNA integrity impacts sequencing read length Read length comparison for samples sheared during the extraction process. Comparison of N50_Q7,_ mean read length_Q7_ and median read length_Q7_ between untreated samples (#10 and #27) and the DNA sample sheared during DNA extraction as shown in Figure 2 #3 (#9).

| Sample | Size selection | N50<sub>Q7</sub> [kb] | Mean<sub>Q7</sub> [kb] | Median<sub>Q7</sub> [kb] | Yield [Gb] | Yield<sub>Q7</sub> [Gb] |
| --- | --- | --- | --- | --- | --- | --- |
| 10 | NO | 25.8 | 12.4 | 6.2 | 6.0 | 5.9 |
| 27 | NO | 26 | 13.2 | 7.5 | 7.8 | 7.4 |
| 9 | sheared during extraction | 9.2 | 4.9 | 2.5 | 3.5 | 3.5 |

**Table 3.** Targeted mechanical DNA shearing does not increase sequencing throughput. Read length comparisons for unsheared and sheared samples. Comparison of N50_Q7,_ mean read length_Q7_ and median read length_Q7_ of untreated samples (#10 and #27) and sheared (g-covaris tube) samples (#4 and #23) (Figure 3).

| Sample | Size selection | N50<sub>Q7</sub> [kb] | Mean<sub>Q7</sub> [kb] | Median<sub>Q7</sub> [kb] | Yield [Gb] | Yield<sub>Q7</sub> [Gb] |
| --- | --- | --- | --- | --- | --- | --- |
| 10 | NO | 25.8 | 12.4 | 6.2 | 6.0 | 5.9 |
| 27 | NO | 26 | 13.2 | 7.5 | 7.8 | 7.4 |
| 4 | g-covaris | 18.4 | 11.8 | 9.5 | 4.8 | 4.7 |
| 23 | g-covaris | 17.9 | 11.2 | 8.5 | 7.2 | 7.0 |

**Table 4.**
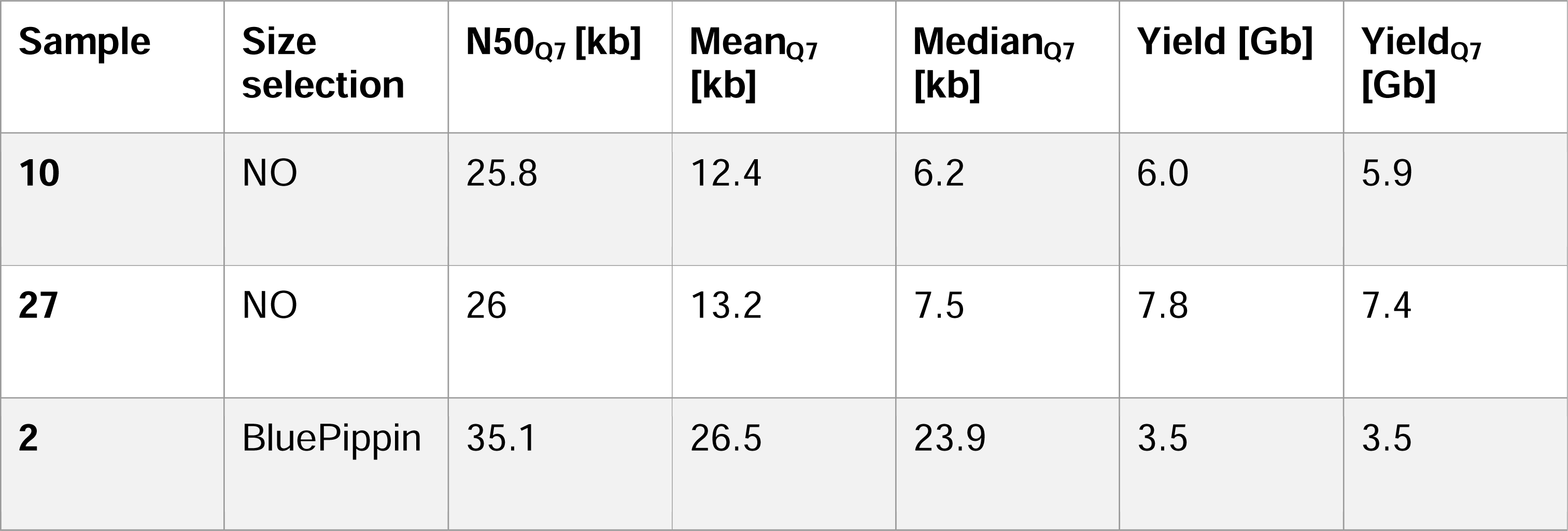
High-pass size selection increases read length statistics. Read-length comparisons for BluePippin size-selected samples. Comparison of N50_Q7,_ mean read-length_Q7_ and median read-length_Q7_ of untreated samples (10) and (27) and Blue-Pippin size-selected samples (2) (Figure 3).

**Figure 4.**
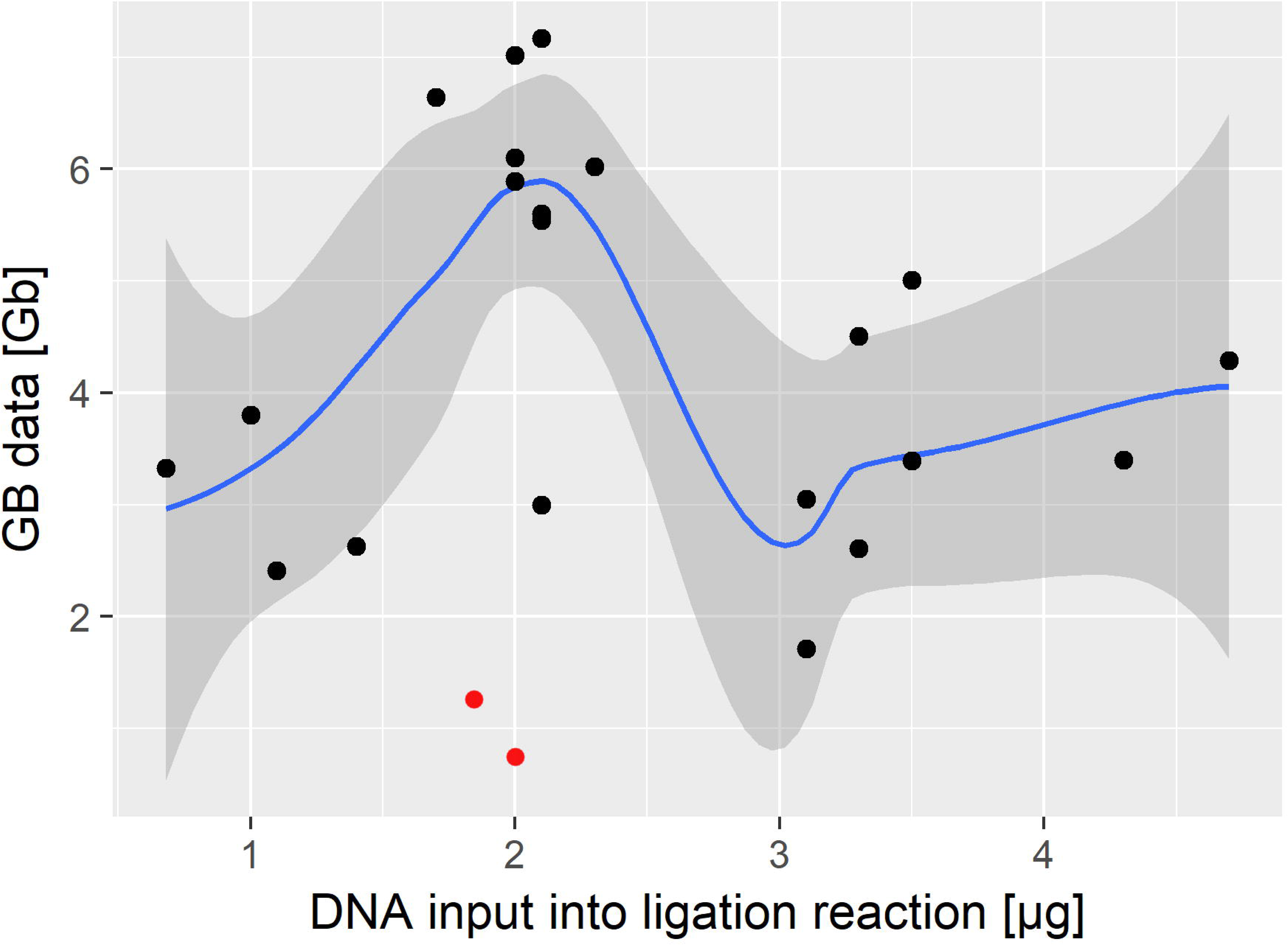
Optimized DNA input into the sequencing adapter ligation reaction. DNA input [µg] into the adapter-ligation reaction of the 1D library preparation (x-axis) versus final sequence yields [Gb]. We achieved highest sequencing throughput by adding approximately 2ug of FFPE and end-repaired DNA into the adapter-ligation step. This optimum was identified empirically and is likely related to the best ratio of free DNA-ends to available DNA sequencing adapters in the ligation reaction. The red points mark outliers with low yields due to a broken flowcell membrane and low yield due bad quality DNA input (Table 1). These points were excluded from the calculation of the smoothed line.

### Altering DNA fragment length and DNA read length

Several factors influence DNA stability during extraction, including chemical properties of the buffer and the physical forces applied during tissue homogenisation, phase separation, and pipetting (Klingstrom, Bongcam-Rudloff, & Pettersson, 2018). The buffer composition is the least flexible factor, especially for difficult tissues such as field samples of eucalyptus leaves that require complex buffers for DNA extraction (see above). In contrast, the conditions during tissue homogenisation can be adjusted more easily by changing treatment type and length. Optimizing these parameters is very important when optimizing DNA fragment length.

To demonstrate this effect, we compared DNA fragment length with sequencing read lengths between two sets of samples that were subjected to different tissue homogenisation procedures. Our standard tissue homogenisation method for eucalyptus leaves consisted of crushing frozen samples for 35 seconds with two 5 mm metal beads in a Qiagen tissuelyser at 24 Hz. To maintain the frozen state, each Eppendorf tube as well as the grinding rack were frozen in liquid nitrogen before the homogenisation step. In an attempt to improve throughput, we tested the effect of homogenizing samples in larger batches, which likely led to a situation where not all samples were completely frozen throughout the procedure. This small change in handling clearly impacted the DNA fragment length distribution as estimated by 0.8% agarose gel electrophoresis. DNA samples extracted using our standard method migrated largely as a single high molecular weight DNA band at the upper limit of resolution (∼23 kb), and well above the 10 kb size standard. For this sample we observed only a light smear visible to 2.5 kb. In contrast, the tissue sample treated in larger batches showed an enhanced low molecular weight smear visible to 1 kb (Figure 2) in addition to the large HMW band. This suggests that the average DNA fragment length was reduced in this sample. To more accurately assess the effect of the change in tissue handling, we ran the second DNA extraction on a single flowcell, and compared the results to those of two flowcells loaded with DNA prepared using the standard (constantly frozen) tissue handling method. The relatively subtle increase in visible DNA smearing on the agarose gel (Figure 2) belied a drastic shift in read length distributions; the mean read length dropped from ∼13 kb to 4.9 kb, and the median from ∼7 kb to 2.5 kb (Table 2). This illustrates that even a slight change in DNA smearing can have a huge impact on sequencing output. It is therefore important to carefully assess DNA fragment length, if possible by comparison to other samples, by agarose gel or PFGE to avoid short sequence reads.

Because our focus for this project was on generating reads >5 kb to assemble a repeat-rich genome *de novo*, we reasoned that it would be beneficial to remove smaller DNA fragments (<1-2 kb) from all samples. AMPure XP beads can be used to size-select DNA fragments in the range 100-500 bp (He, Zhu, & Gu, 2013; Schmitz & Riesner, 2006). However, it is not possible to remove DNA fragments larger than ∼500 bp with AMPure XP beads (Figure 5) as adding less than 0.4 vol (V/V) of bead solution causes the NaCl concentration to fall below 0.4 M, leading to complete sample loss (He et al., 2013). We reasoned that by adjusting the PEG and NaCl concentrations, which precipitate DNA in a cooperative manner, we might be able to select a higher average DNA fragment length, and thereby remove unwanted smaller DNA fragments (Lis & Schleif, 1975; Ramos, de Vries, & Ruggiero Neto, 2005). Using 0.8 vol (V/V) of our adjusted SPRI beads mixture (which translates to final PEG concentrations of 4.8% (v/v) and 0.7 M NaCl) we were able to remove DNA fragments of up to 1.5 kb (Figure 5) (Schalamun & Schwessinger, 2017). We used this adapted SPRI beads mixture subsequently for DNA sample clean up and during library preparation.

**Figure 5.**
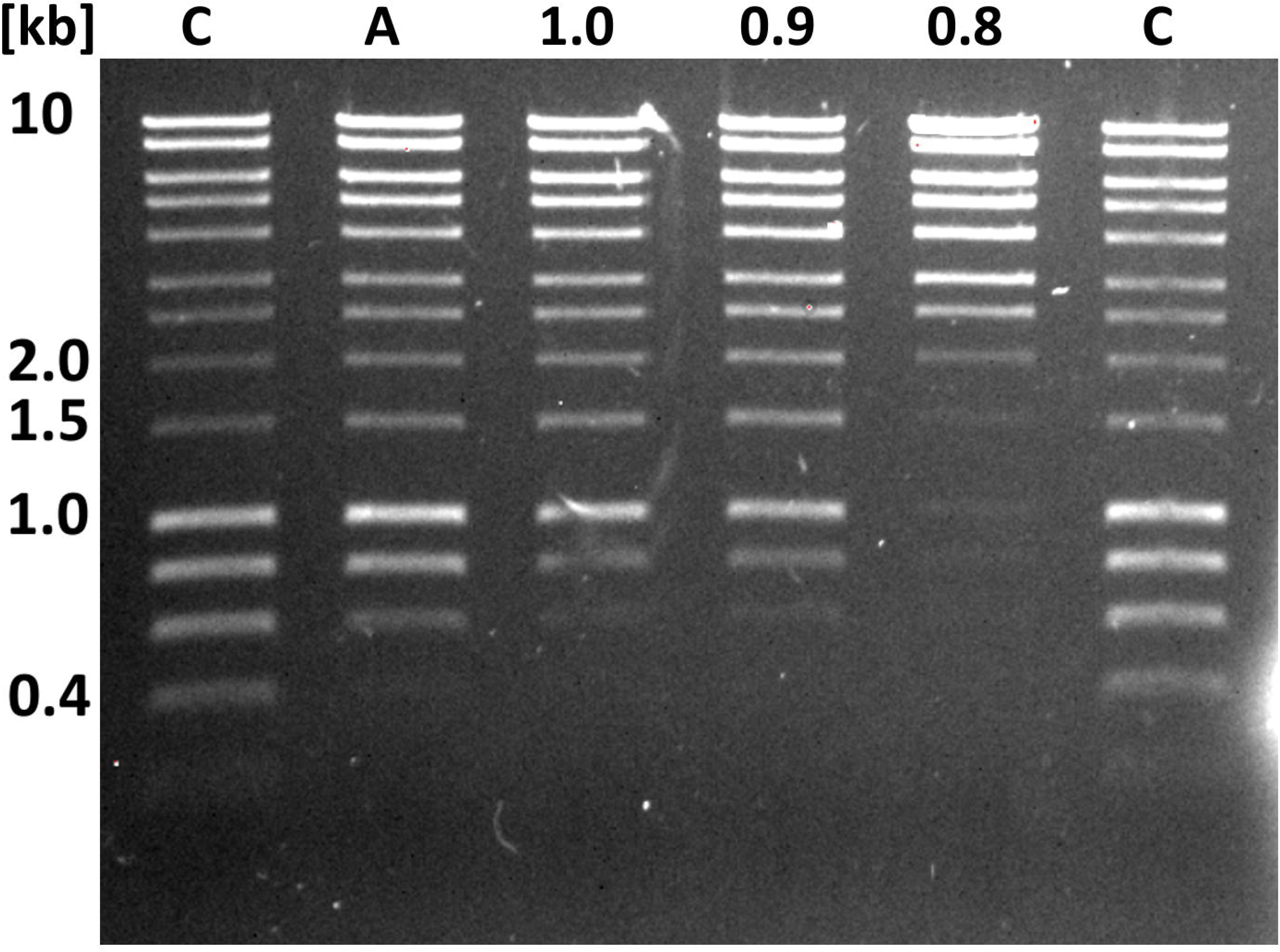
Improved DNA size selection using an adapted PEG-NaCl-SPRI beads protocol. Each lane represents 80 ng DNA before size selection. Lanes C contain the HyperLadder 1 kb (BioLine) as untreated control. Lane A is DNA ladder size selected with 0.45 vol (V/V) Agencourt AMPure XP beads. Lane 1.0, 0.9, and 0.8 are DNA ladder size selected with the adapted PEG-NaCl-SPRI beads solution.

We next assessed the effect of DNA shearing and gel-based size-selection procedures on sequencing throughput and read length distribution. In the case of DNA shearing our hypothesis was that a more unimodal size distribution of shorter DNA fragments with a peak of about 20 kb (Figure 3) would increase sequencing throughput. We used g-TUBEs with a benchtop centrifuge to shear DNA by forcing it through a µm mesh. DNA shearing did not increase yield, but did affect the read length distribution (Table 3). Compared with non-sheared samples, the sequence read length distribution from sheared reads shifted to smaller values and peaked at about 11 kb (Figure 6), with an N50_Q7_ of 18 kb, compared to an N50_Q7_ of ∼26 kb from the unsheared samples (Table 3). Whereas Q7 presents the default quality threshold from the basecaller. Interestingly, the median read length from the sheared DNA samples increased to 7.5 kb from 6.5 kb when compared to unsheared DNA. At the same time low quality short reads were reduced in the sheared samples. Possibly, removing long DNA fragments (> 50 kb) leads to fewer low quality reads caused by long DNA molecules being stuck in the pore, at least when using the R9.5 pore chemistry. This highlights that filtering reads based on their q-scores, as well as removing short sequencing reads, may help to avoid error propagation during downstream analyses of the data.

**Figure 6.**
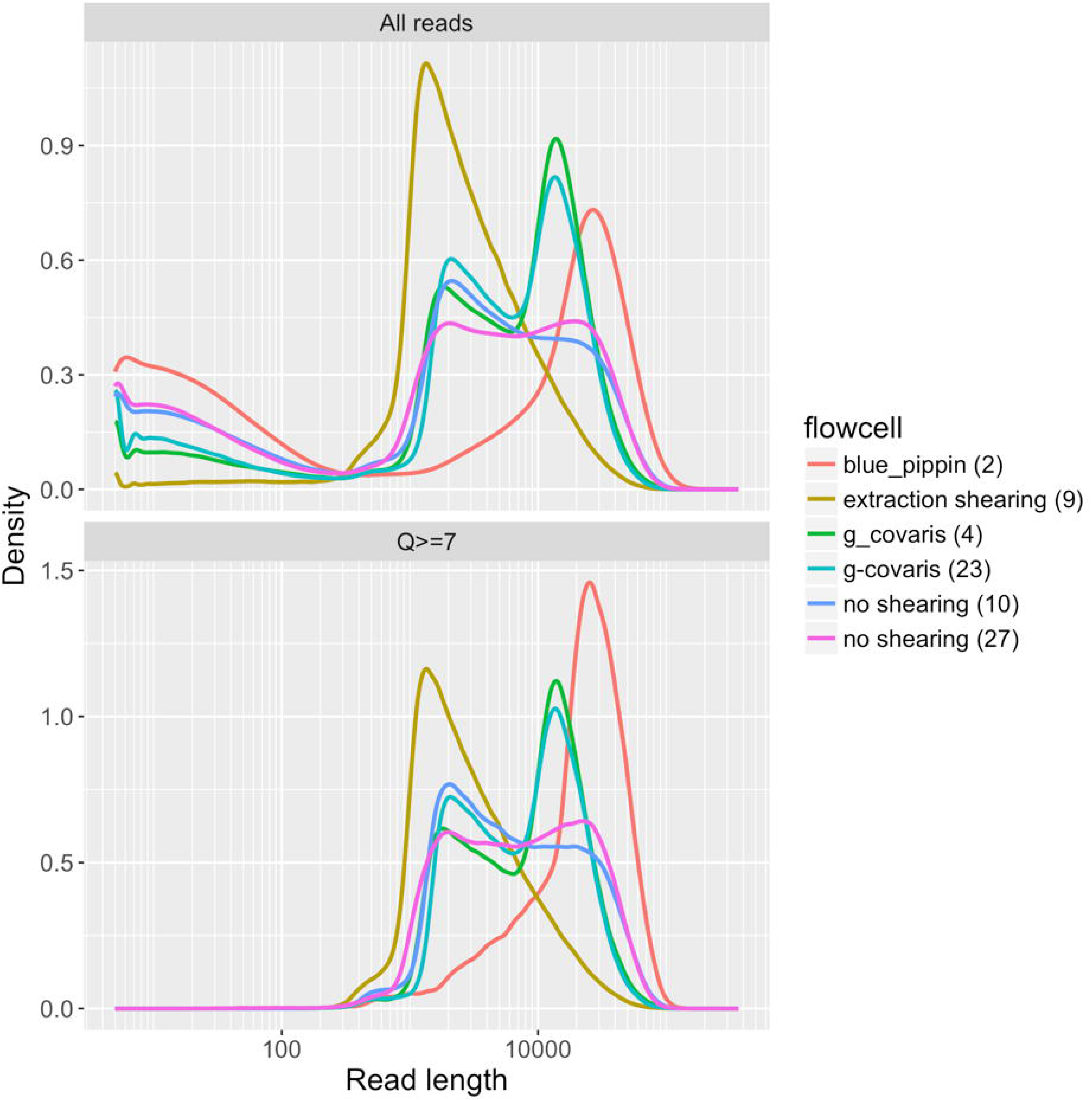
The impact of DNA extraction protocol on the distribution of read lengths from ONT sequencing. Each line represents the read length distribution for a single flowcell. The x-axis shows the read lengths on a log scale, and the y-axis shows the density of reads at a particular length. The top panel shows data for all reads, and the bottom panel shows the same data, but with reads that have a mean quality (Q) score less than 7 removed.

**Figure 7.**
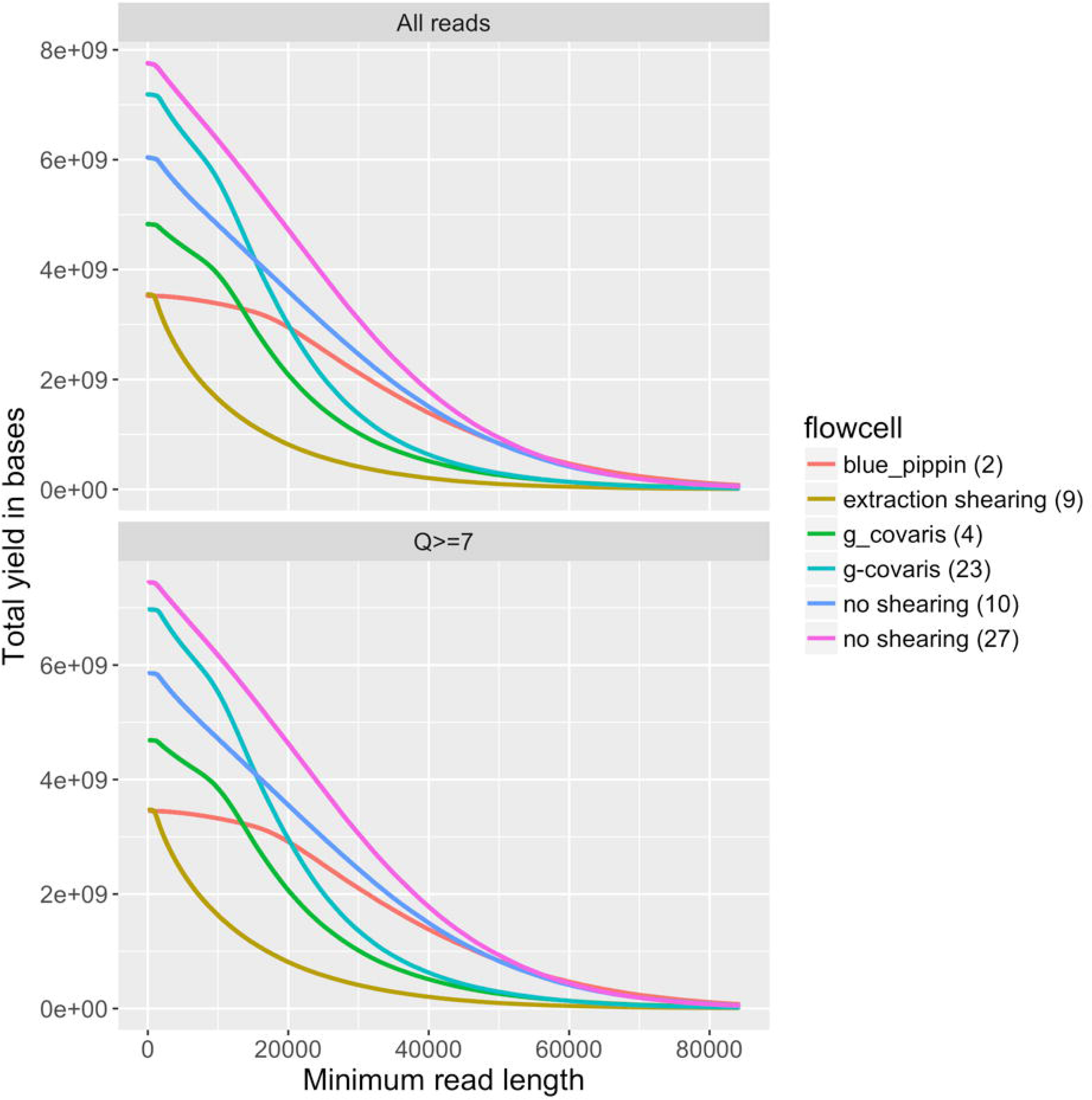
The impact of DNA extraction protocol on the yield of ONT sequencing. Each line represents a single flowcell. The y axis shows the yield of in bases, and the x axis shows the minimum readlength at which the yield was calculated. For example, the yield of reads longer than 20KB from each flowcell can be compared by comparing the height of the lines at the 20KB point on the x axis. The plot shows that while using the Blue Pippen improved the distribution of read lengths (insofar as the red line is relatively flat below 20KB, showing that only a small proportion of the sequenced bases were in reads shorter than 20KB), we were able to obtain higher yields of reads >20KB from the libraries that were prepared without the Blue Pippen (blue and pink lines, labelled ‘no shearing’). These flowcells also contained a considerable yield of reads between 1KB and 20KB, which may be useful for many applications.

We also tested the effect of removing DNA fragments below 20 kb by size selection using the BluePippin system in the high-pass mode which enables the collection of DNA molecules above a certain size. When we applied the 20 kb high-pass filter we were able to remove DNA fragments less than 20 kb while maintaining the high molecular weight size distribution (Figure 3). After sequencing, the read length N50_Q7_ increased to 35 kb from 26 kb, while the mean and median read-lengths increased to 26 and 23 kb from 12 and 6.5 kb respectively (Table 4 and Figure 3). The main drawbacks of BluePippin high-pass size selection were the high sample loss (65-75%), the increase in cost, and prolonged sample handling.

Overall, we did not employ DNA shearing using g-TUBEs or size selection via BluePippin in our final sequencing protocol even though they improved sequencing read-length distributions. In our case, the high quality sequencing results achieved with our standard protocol using the improved SPRI beads mixture did not warrant the additional time and financial investment required when incorporating g-TUBEs DNA shearing or BluePippin size selection into our workflow.

### Real time and between run evaluation

The software MinKNOW makes it possible to perform a real time monitoring during the MinION sequencing run. Interpreting the pore signal statistics and the length graph during the first two hours of sequencing gives the user a clear idea if the run should be continued or stopped. We used this feature of MinKNOW to optimize our runs. First, we evaluated pore occupancy, defined as the ratio of ‘in strand’ (light green) to the sum of ‘in strand’ plus ‘single pores’, after one hour. A high pore occupancy (>70%) indicates successful library preparation and is predictive of a high final sequencing output. If the pore occupancy was below 70% we stopped the run, washed the flowcell and loaded a new library to ensure high throughput per flowcell (Figure 8). We reasoned that these low throughput runs were usually due to insufficient DNA molecules being ligated to sequencing adapters during the library preparation. We found that we had to load at least 1 µg library DNA onto a flowcell to achieve acceptable yields (Figure 4). To ensure sufficient adapter ligated DNA, we started library preparation with at least 4 µg of high quality starting DNA to account for potential losses during the SPRI bead clean-up steps. A second pore statistic to consider is the number of unavailable pores, e.g. ‘zero’ (black), ‘unavailable’ (light blue), or ‘active feedback’ (pink) (Mayjonade, 2018). If these numbers increase too quickly in the first few hours of the run it is likely that the library is contaminated and the pores are being irreversibly blocked or damaged, or that the membrane has ruptured. If recognized early enough the flowcell can be washed and a new library loaded, but the pores cannot always be recovered. Furthermore, the length distribution from the length graph can be easily assessed and, if unsatisfactory, the library exchanged for a separately prepared sample (Figure 8). We also recommend to track the sequencing run with a continuous screenshot application (e.g. newlapse for linux, https://github.com/mtib/newlapse), in addition to visual inspection during the first few hours of the sequencing run. This enables continuous monitoring and assessment of unusual sequencing behaviours out of hours.

**Figure 8.**
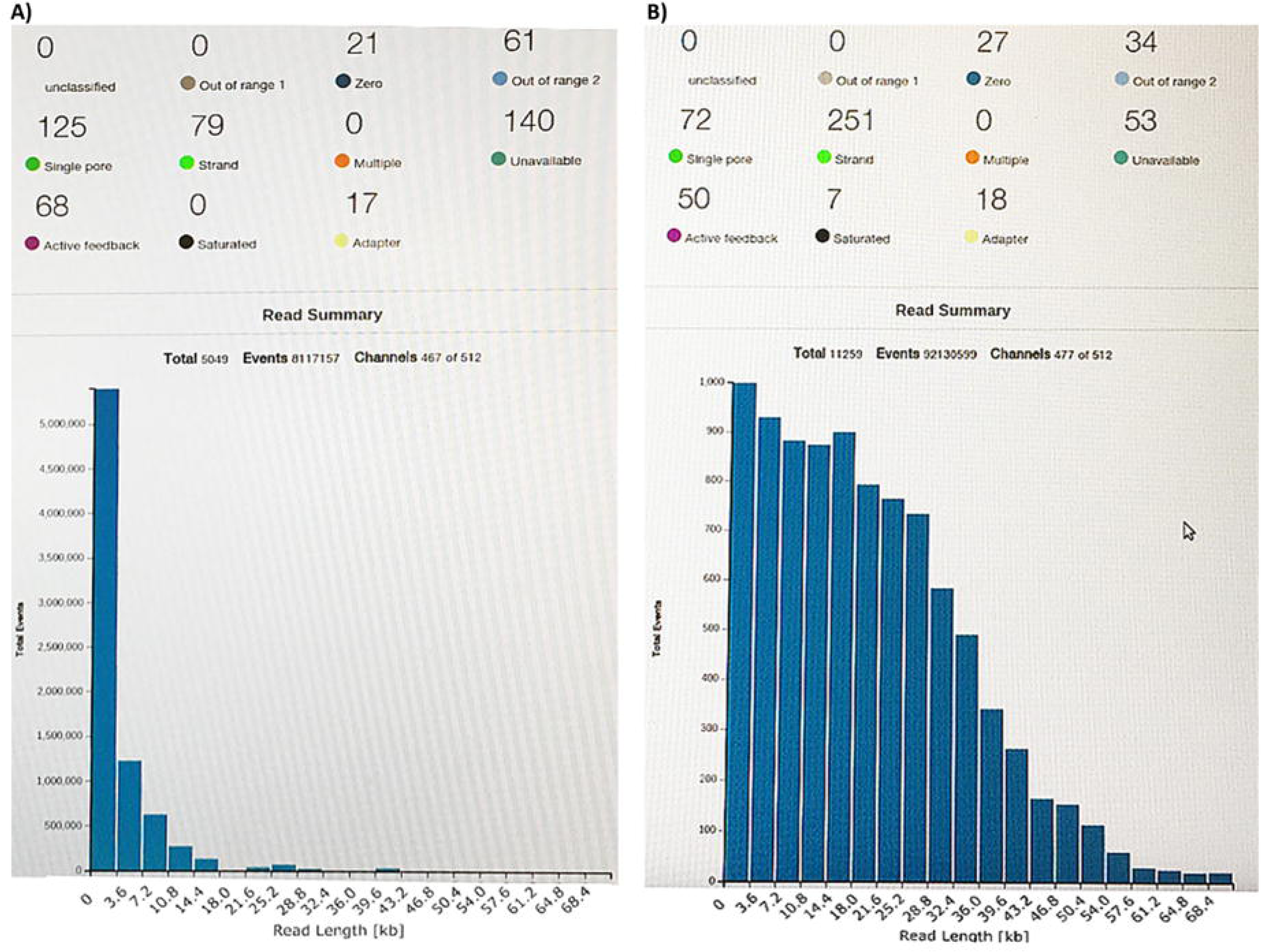
Real time analysis of sequencing runs via the MinKNOW graphical user interface. Both panels (A and B) show the MinKNOW interface two hours into a run. Panel A illustrates an unsatisfactory sequencing run where read length is short, pore occupancy poor (∼40%) and many pores are not available for sequencing any more (see main text for details). This run was aborted after two hours to not to waste this flow cell and to reload an improved library. Panel B illustrates a satisfactory sequencing run with excellent read length distribution, good pore occupancy (∼80%), and most pores still readily available for sequencing.

**Figure 9.**
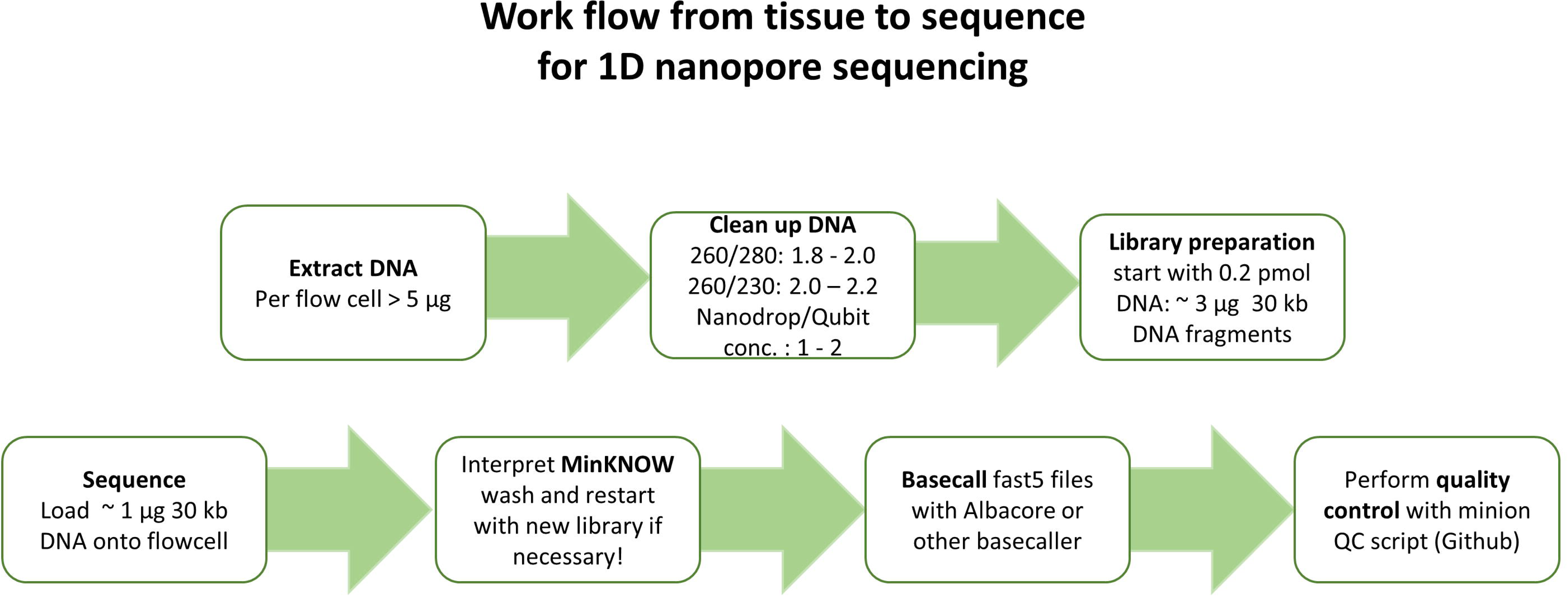
MinION Nanopore sequencing workflow to optimize sequencing output.

One key to ongoing optimisation of flowcells in our laboratories was the tracking of all parameters for each sequencing run using our monitoring spreadsheet (Supplemental Table 2) and a continued comparison of the output of each additional flowcell. After running each flowcell, we used ONT’s Albacore 2.0 basecaller to convert the raw signal data from the MinION into DNA sequence data in fastq format. Albacore 2.0 produces a sequencing_summary.txt file which contains a summary of every sequencing read, and can be used for rapid assessment of each flowcell using the minionQC script (https://github.com/roblanf/minion_qc). After basecalling each flowcell, we ran this script and examined in detail the length and mean quality distributions of the reads, and the physical performance map of the flowcell. This allowed us to continually evaluate and improve our protocols for each flowcell. Before we were halfway through our project, we were able to reliably and repeatedly obtain more than 6 Gb of data from each flowcell, with mean read lengths consistently above 12 kb.

## Discussion

Here we present a complete workflow to establish MinION long read sequencing in any laboratory. We highlight the importance of clean high molecular weight DNA for successful sequencing runs and provide detailed wet lab DNA extraction and purification protocols that include size selection. All these protocols and many others applicable to different starting material, some provided by other community members, are freely available on the open-access protocol sharing repository protocols.io in form of a MinION user group (https://www.protocols.io/groups/minion-user-group-with-fungi-and-plants-on-their-mind) (Schwessinger, 2016). We encourage others to contribute to this open science platform to accelerate research and for the community to save costs when establishing long read DNA sequencing in their own laboratory. High quality ‘living’ protocols with careful run and run-to-run evaluations as described here will facilitate knowledge generation instead of constant ‘reinvention of the wheel’.

## Acknowledgment

We would like to acknowledge fruitful discussion, leading to and improving this manuscript, with the following; Louise Judd, Ken McGrath, Baptiste Mayjonade, David Hayward, Josh Quick, and Megan McDonald. We would also like to acknowledge all contributors of the MinION usergroup on protocols.io for sharing their protocols openly.

## Methods

### Tissue collection

*Eucalyptus pauciflora* leaf tissue was collected from Thredbo, New South Wales (NSW), Australia. After harvesting the young twigs were transported in plastic bags and stored in darkness at 4°C in water until DNA extraction.

### High molecular weight DNA extraction and clean up

We extracted high molecular weight DNA based on Mayjonade’s DNA extraction protocol optimized for our eucalyptus samples (Mayjonade et al., 2016). Each extraction was carried out with 800 - 1000 mg leaf tissue which was cut into small pieces and split between 8 separate 2 mL Eppendorf tubes, each containing 2 metal beads of 5 mm in diameter, before freezing in liquid nitrogen. We lysed the tissue mechanically by grinding using the Qiagen tissue lyzer II for 35 seconds at 25 Hz. Pulverised tissue was suspended in 700 µL SDS lysis buffer (1% w/v PVP40, 1% w/v PVP10, 500 mM NaCl, 100 mM Tris-HCl pH 8.0, 50 mM EDTA, 1.25% w/v SDS, 1% w/v sodium metabisulfite, 5 mM DTT, Milli-Q water and heated to 64°C for 30 minutes to inactivated DNases. One µL RNase A (10 mg/mL) (Thermo Fisher) per 1 mL lysis buffer was added to the mixture after the heat treatment, followed by incubation at 37°C for 50 minutes at 400 rpm on a thermomixer. Twenty minutes into the incubation we added 10 µl Proteinase K (800 Units/mL) (NEB). To precipitate proteins, the tubes were cooled on ice for 2 min before adding 0.3 vol (210 µL) 5 M potassium acetate pH 7.5 and mixed by inverting the tube 20 times. The precipitates containing leaf material and proteins were removed by centrifugation at 8000 *g* for 12 min at 4°C. We transferred the supernatants to new tubes and purified the DNA from solution as described below in “Removal of small DNA fragments < 1.5 kb with optimized SPRI beads”.

We further purified the samples using a chloroform:isoamylalcohol extraction. The eight aqueous DNA solutions were pooled to a total of 400 µL to which one volume of chloroform:isoamylalcohol was added, mixing by inversion for 5 minutes. The phases were separated by centrifugation at 5000 *g* for 2 minutes at room temperature (RT). We transferred the upper, DNA containing phase to a fresh tube performing another round of the chloroform:isoamylalcohol purification. After the second extraction the DNA was precipitated by adding 0.1 volume 3 M sodium acetate pH 5.3 and 1 volume 100% cold ethanol, followed by centrifugation at 5000 *g* for 2 min at RT. The short centrifugation at low speed may reduce DNA yields but potentially precipitates longer fragments in favor of shorter fragments. The transparent pellet was washed with 70% ethanol and resuspended in 50 µL 10 mM Tris-HCl pH 8.0 for 2 h at room temperature. The solubilised DNA was stored at 4°C until library preparation, for a maximum of 10 days.

### DNA size selection

#### g-TUBE shearing

We processed 5 µg of pure HMW DNA through a g-TUBE (Covaris) in an Eppendorf 5418 centrifuge at 3800 rpm for a total of 2 minutes.

#### BluePippin size selection

We used the BluePippin system (Sage science) with 0.75% dye-free Agarose cassettes and S1 marker, selecting for fragments > 20 kb using 6 µg sample for each lane following the manufacturer’s instructions.

### Removal of small DNA fragments < 1.5 kb with optimized SPRI beads

In order to purify and remove small fragments from our DNA samples we optimized a SPRI beads solution which we used for clean ups and library preparations. The improved beads solution consists of 11% PEG 8000, 1.6 M NaCl, 10 mM Tris-HCl pH 9.0, 1 mM EDTA, 0.4% Sera-Mag SpeedBeads (GE Healthcare PN 65152105050250) (Schalamun & Schwessinger, 2017). For the clean up procedure, 0.8 vol of this beads solution was mixed with the DNA sample and incubated on a hula mixer for 10 min. After a brief (pulse) centrifugation step in a microcentrifuge, we placed the tube in a magnetic stand so that the beads bound to the rear of the tube, allowing for removal of the supernatant. We then washed the beads twice with 1 mL 70% ethanol, keeping the tube on the magnetic stand throughout the wash procedure to avoid loss of DNA bound to the beads (the tube can be rotated 360° within the stand, allowing comprehensive washing while ensuring bead retention). After the second wash we centrifuged the tube briefly again to remove the last traces of ethanol. The beads were dried for no longer than 30 s before elution of the DNA in 50 µL Tris-HCl pH 8.0 preheated to 50°C, for 10 min.

### DNA Quality control

DNA concentrations were determined using the Qubit dsDNA BR (Broad Range) assay kit (ThermoFisher). The purity of the sample was measured with the NanoDrop, assessing curve shape, the 260/280 nm and 260/230 nm values, and congruence of concentrations with the Qubit values. The DNA was examine after 0.8 % agarose gel electrophoresis containing 0.001% (v/v) SYBR Safe dye (ThermoFisher) in 1X TBE buffer (10.8 g/L Tris base, 5.5 g/L boric acid, 0.75 g/L EDTA, pH 8.3) for 45 minutes at 100 V. For higher resolution, pulsed field gel electrophoresis (PFGE) was used with a 1.5% agarose gel in 0.5X TBE running buffer, run for 17.7 hours at 6 V/cm and 1.4 s initial and 13.5 s final switch time. The gel was stained after the electrophoresis with 5 µL SYBR Safe dye in approximately 200 mL Milli-Q water.

### Library preparation and sequencing

We followed the 1D ligation protocol SQK-LSK108 selecting for long reads but instead of the recommended AMPure XP beads we used our optimized SPRI beads solution (Schalamun, 2017). We started the library preparation with 4 µg HMW DNA and used 1 µg of the resultant library DNA for sequencing on a R9.5 flowcell. The sequencing software MinKNOW version 1.7.3 was installed on a computer with minimum of 4 cores running a Linux operating system (Ubuntu 14.4).

## Supplemental material

**Supplemental Table 1. Raw data of all quality control variables tracked during DNA extraction, library preparation, and sequencing run summary statistics described in this study.**

**Supplemental Table 2. Template spreadsheet to use for tracking quality control variables for DNA extraction, library preparation, and sequencing run summary statistics.**

**Supplemental Figure 1: Sequencing yield depends on quality control statistics** Comparison of sequencing yield (as GB of called data) versus all measured QC statistics. Positive correlations that are not directly related to yield are seen with sample number, the number of channels seen during flow cell QC, and the measured initial input library concentration. Lines indicate fitted quantile regression lines at 25%, 50% and 75% (i.e. lower quartile, median, and upper quartile respectively). The script for plotting the quality control statistics vs sequencing yield is provided on github (https://github.com/gringer/minion-user-group).

