## Supplementary figures and images for "A comprehensive toolkit to enable MinION sequencing in any laboratory"

### Supplementary Materials

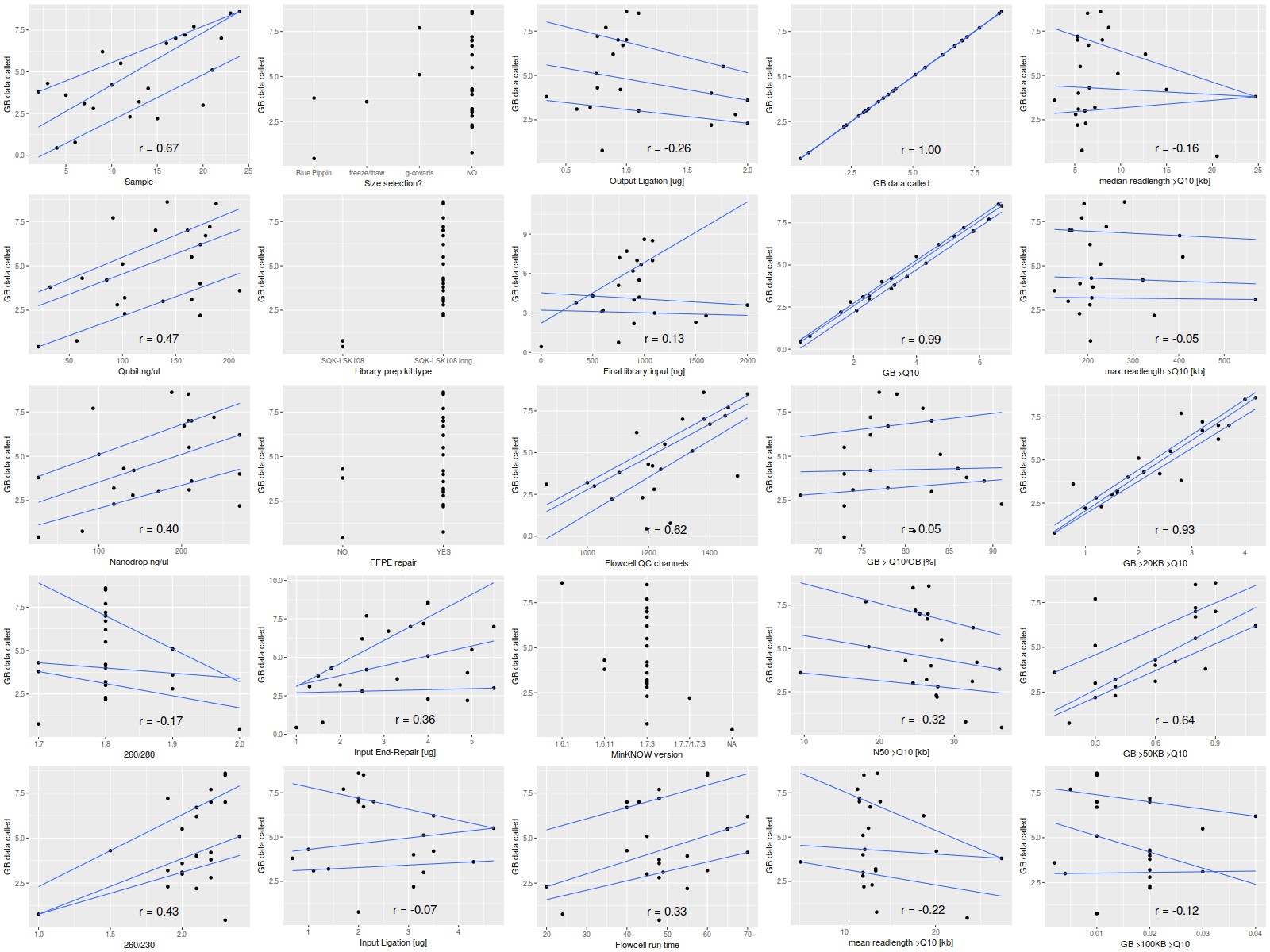
